# Stable motifs delay species loss in simulated food webs

**DOI:** 10.1101/2021.04.06.438635

**Authors:** Alyssa R. Cirtwill, Kate Wootton

## Abstract

Some three-species motifs (unique patterns of interactions between three species) are both more stable when modeled in isolation and over-represented in empirical food webs. This suggests that these motifs may reduce extinction risk for species participating in them, ultimately stabilizing the food web as a whole. We test whether a species’ time to extinction following a perturbation is related to its participation in stable and unstable motifs and assess how motif roles co-vary with a species’ degree or trophic level. We found that species’ motif roles are related to their times to extinction following a disturbance. Specifically, participating in many omnivory motifs (whether in absolute terms, as a proportion of the species’ role, or relative to other species in the network) was associated with more rapid extinction, even though omnivory has previously been identified as a stable motif. Participating in the other three stable motifs (three-species chain, apparent competition, and direct competition) was associated with longer times to extinction. While motif roles were associated with extinction risk, they also varied strongly with degree and trophic level. This means that these simpler measures of a species’ role may be sufficient to roughly predict which species are most vulnerable to disturbance, but the additional information encapsulated in a motif role can further refine predictions of vulnerability. Moreover, where researchers are *a priori* interested in motif roles, our results confirm that these roles can be interpreted with respect to extinction risk.

## Introduction

The connections between food-web structure and the extinction risk of species within the food web have interested ecologists since at least the 1970’s (May, 1972). After early findings suggested that a large, randomly-connected network is unlikely to retain all species after small perturbations (Gardner & Ashby, 1970; May, 1972), non-random structures that might stabilize food webs (allowing all species to persist) were quickly identified. These structural features include nestedness (Allesina & Tang, 2012; Sauve *et al.*, 2014), modularity (Sauve *et al.*, 2014; Thébault & Fontaine, 2010) and skewed distributions of link strengths (McCann *et al.*, 1998; Gross *et al.*, 2009; Rooney & McCann, 2012; Wootton & Stouffer, 2016a).

Although important, these global-scale properties (i.e. properties of the network as a whole, see Fig. 1) can mask important differences in network structure (Simmons *et al.*, 2019) and do not provide information about differences between species within the same network (Cirtwill *et al.*, 2018). Two networks with the same connectance and similar values of nestedness and modularity may still have quite different meso-scale (i.e., more detailed than global-scale) arrangements of links. These meso-scale structures, described by frequencies of *motifs* (unique patterns of *n* interacting species), define the local neighborhood of a focal species and reflect its direct and close indirect interactions (Fig. 1).

**Figure 1:**
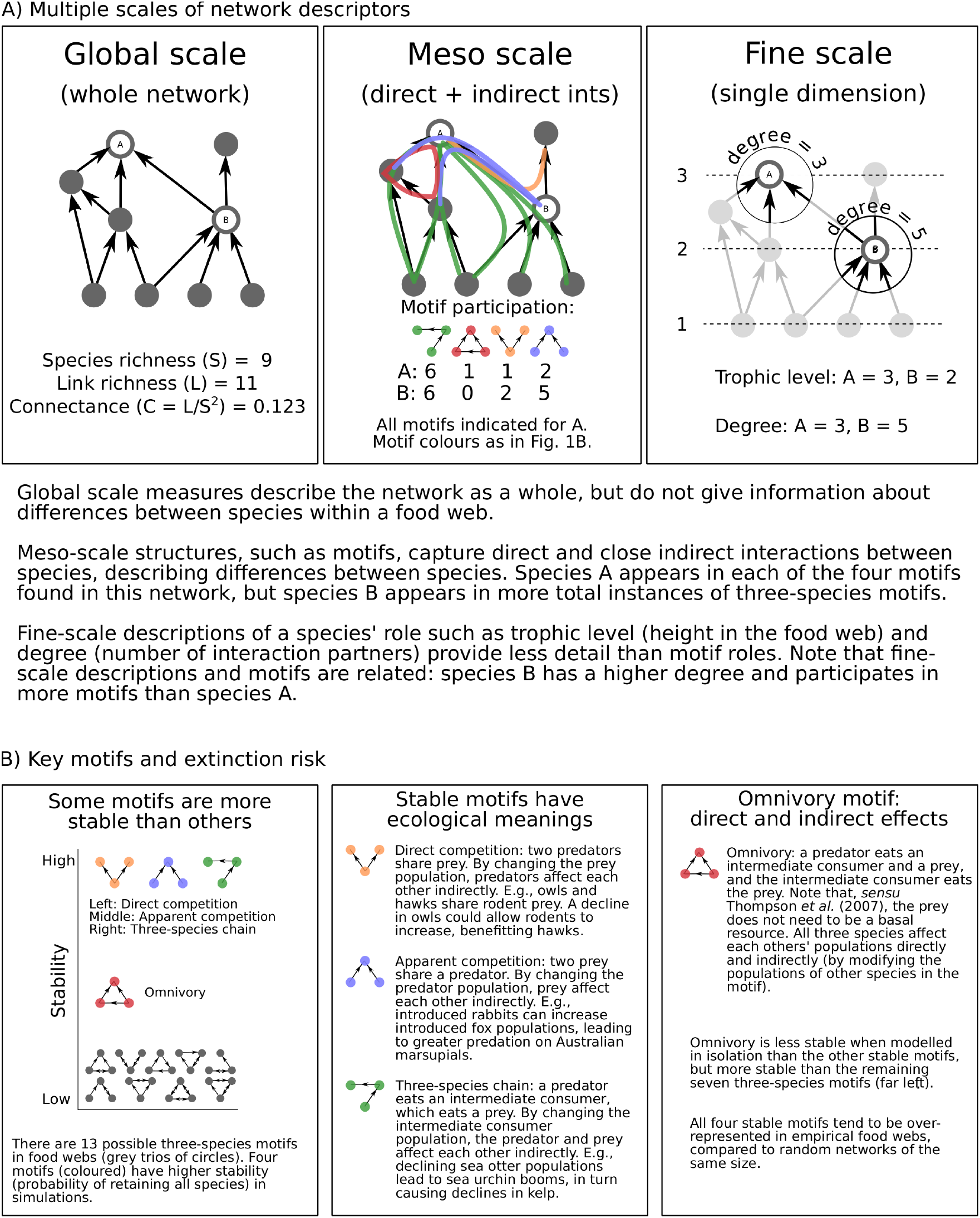
A brief introduction to motifs.

In addition to providing species-level information on network structure, there are early indications that some meso-scale structures may tend to stabilize food webs (Prill *et al.*, 2005; Borrelli *et al.*, 2015; Monteiro & Faria, 2016). This possibility is supported by the fact that empirical food webs tend to contain more *three-species chains* and either more *omnivory* (*sensu* Thompson *et al.*, 2007) motifs or more *apparent competition* and *direct competition* motifs than random networks (Stouffer *et al.*, 2007). These four motifs make up, on average, about 95% of all three-species motifs in food webs (Stouffer & Bascompte, 2010). The high frequencies of these four motifs in observed networks suggest that they may be beneficial to the network containing them or, in other words, that more stable motifs appear more frequently in empirical food webs because unstable motifs are more likely to disappear as the species within them go extinct (Borrelli *et al.*, 2015; Borrelli, 2015) or links between species are lost (Tylianakis *et al.*, 2010).

When modeled in isolation, three species arranged in *three-species chain, apparent competition, direct competition*, and *omnivory* motifs are more likely to all persist than trios of species arranged in other motifs (Borrelli, 2015). By damping perturbations and maintaining more constant populations of the species participating in them, these ‘stable’ motifs may contribute to the stability of the network as a whole (Borrelli, 2015). The greater chances of maintaining all species in a stable motif could explain the over-representation of these motifs in empirical food webs if species and interactions in less-stable motifs are more likely to be “pruned” from empirical communities over time. We expect that stable structures will endure for longer than unstable ones, and so it is not surprising that stable structures should be over-represented in empirical communities (Borrelli *et al.*, 2015).

Given these early indications that some motifs are more stable than others, we may also expect that a species’ role– here defined as the frequency with which it participates in different motifs –could affect its probability of extinction following a perturbation.

Specifically, we expect that species participating more frequently in the stable motifs identified by Borrelli (2015) are less likely to go extinct than species whose roles are dominated by other motifs. Testing this hypothesis is complicated by the fact that species’ motif roles are not independent of other aspects of fine-scale network structure. In particular, a species’ degree (number of predators and prey) is likely to strongly affect its motif role. The more interaction partners a species has, the more motifs in can participate in (Fig. 1). This is likely to lead species with higher degrees to participate in a wider variety of motifs as well as participate more frequently in the four common, stable motifs. Species with high in-degrees (more prey) are also generally less likely to go extinct, as are species at lower trophic levels (Cirtwill *et al.*, 2018). Any test for a relationship between motif roles and time to extinction should therefore take degree and trophic level into account to avoid confounding these effects.

Here, we investigate the relationship between species roles and extinction risk by simulating the removal of species from simulated networks at stable equilibria. We test 1) whether species’ roles are related to their time to extinction following the removal of another species in the network, 2) whether participation in particular motifs (especially the stable motifs described above) is correlated with time to extinction, and 3) whether these correlations are driven by potential relationships between species’ participation in various motifs and simpler definitions of species roles (degree, trophic level). Taken together, these tests show that species are generally consistent in their vulnerability to disturbances, regardless of the location in the network of that disturbance, and this vulnerability is shaped by both motif roles and other network parameters.

## Methods

### Generating networks and extinctions

#### Generating food webs

We simulated a suite of food webs based on the probabilistic niche model, which assigns predator-prey links based on the body-mass ratios between individuals of different species (Williams & Martinez, 2000; Delmas *et al.*, 2017). The meso-scale structure of niche-model networks closely mimics that of empirical food webs (Stouffer *et al.*, 2007). To ensure that we captured a variety of realistic community sizes and structures, we generated networks ranging between 50 and 100 species (in steps of 10) with connectance values between 0.02 and 0.2 (in steps of 0.02). The range of network sizes was chosen to reflect moderately well-sampled empirical webs while working within our computational limits, while the range of connectance values was chosen to cover that observed in most empirical food webs (Dunne *et al.*, 2002). We generated a total of 100 networks with each combination of parameters, for a total of 6000 networks. All networks were generated using the function “nichemodel” within the Julia language package *BioEnergeticFoodWebs* (Delmas *et al.*, 2019, 2017). If a simulated network contained any disconnected species (species without predators or prey) or disconnected components (a group of species connected among themselves but not to the rest of the network), the network was rejected and a new network simulated. Finally, networks where the path lengths between each species and a basal resource could not be resolved (i.e., trophic levels were undefined) were rejected and new networks simulated.

After generating the network structure, we simulated community dynamics using the function “simulate” from the Julia language package *BioEnergeticFoodWebs* (Delmas *et al.*, 2019, 2017). This function uses Lotka-Volterra predator-prey models including density dependence and type 2 functional responses for all species (please see Delmas *et al.* (2017) for full details). All non-basal species were designated as vertebrates to ensure a good match between metabolic and predator-prey body-mass ratio values. Metabolic rates in the Lotka-Volterra model are based on each species’ body mass (i.e. mass of a single individual). We assigned relative body masses based on each species’ trophic level, which was, in turn, calculated based on the food-web structure provided by the niche model. After basal species were assigned a body mass of 1, we used a predator-prey body-mass ratio of 3.065 to calculate the relative body masses of higher trophic levels. We selected this ratio based on the estimate for vertebrates (averaged across ecosystem and metabolic types) in Brose *et al.* (2006). We excluded reported body-mass ratios for invertebrates as these could include parasites and parasitoids, which are generally smaller than their prey, and because interactions among vertebrates are better represented in the food-web literature than interactions involving invertebrates.

The persistence of each species in our simulated networks also depends on its population biomass. We randomly assigned initial population biomasses (i.e. cumulative biomass across all individuals of a species) for each species from a uniform distribution [0,1]. Note that population biomasses and individual body masses are not calculated on the same scale. We then simulated community dynamics for 1000 time steps to obtain an equilibrium community. To ensure that species did not ‘recover’ from unrealistically low biomasses during the simulation, we considered a species extinct if it dropped below an arbitrary threshold biomass of 1×10^−5^. When simulating initial (i.e. pre-perturbation or equilibrium) dynamics, we rejected any network where one or more species dropped below this biomass threshold. Consumers were assumed to have no preferences such that the consumption rate *w*_*ij*_ of predator *i* eating prey *j* is equal to 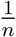, where *n* is the number of prey for predator *i*. If the network did not reach an equilibrium with all species persisting for 1000 time steps, a new set of initial population biomasses was applied and the simulation repeated until an equilibrium with all species persisting was obtained. If a stable equilibrium still had not been reached after 100 sets of randomly-assigned initial biomasses, we discarded the network and simulated another to replace it.

#### Calculating species roles

We were interested in whether species’ roles at equilibrium are related to their response to a perturbation, in this case the removal of another species in the network. We defined each species’ role as the number of times it appears in each unique three-species motif, following Stouffer *et al.* (2012); Cirtwill & Stouffer (2015). Note that each set of three interacting species forms exactly one motif (Cirtwill *et al.*, 2018). As our focus is on how different motifs might affect extinction risk, we do not consider the different positions species may take within a motif. We expect that appearing more frequently in stable motifs (three-species chain, apparent and direct competition, and omnivory) will correlate with lower extinction risk while appearing more frequently in unstable motifs (those containing two- or three-species loops) will be associated with higher extinction risk. Note that cannibalistic links were ignored when calculating motif frequencies within a network and species’ roles, although they were included when calculating connectance. As well as these ‘raw’ motif roles, we calculated ‘degree-normalized’ motif roles for each species by dividing the number of appearances in each motif by the total number of times the species appears in any motif (as in Cirtwill & Stouffer (2015); this total is expected to strongly correlate with degree). Finally, we also calculated ‘network-normalized’ motif roles, defined as the z-score of the number of times a focal species appears in each motif compared to the number of times all species in the network appear in the focal motif. The degree normalization allows us to test whether trends in stability with motif participation are due to differences in the total number of motifs a species appears in, while the network normalization allows us to test whether trends in stability with participation in different motifs are related to how unusual each species is within its community context.

#### Perturbing networks

After identifying species’ roles in the equilibrium networks, we perturbed the networks by removing a single species. After this removal, community dynamics were simulated for 50 rounds of 10 time-steps (500 time-steps total). After each round, any species with a biomass below our threshold of 1×10^−5^ was considered to have gone extinct and its biomass was set to 0. We recorded the biomass of each species after each round, as well as the round in which any additional extinctions occurred. After 500 time-steps, we reset the network to its original state (including all species). We then removed a new species and again simulated community dynamics. We repeated this process until all species had served as the initial removal. We then calculated the mean time to extinction across all removals as an overall measure of each species’ vulnerability (*Appendix S1*). Time to extinction was highly correlated across removals in all combinations of S and C, indicating that this is a robust measure.

### Testing effects of species’ roles on time to extinction

#### Overall motif roles

To test for a relationship between time to extinction and species’ overall roles, we fit a series of PERMANOVAs (Anderson, 2001) relating Bray-Curtis dissimilarity (Baker *et al.*, 2015; Cirtwill & Stouffer, 2015) in species’ raw motif roles to differences in mean time to extinction (Table 1). Due to computational limits, we were unable to fit a single PERMANOVA for all networks. To avoid effects of network size and connectance on time to extinction, we therefore fit separate PERMANOVAs for each combination of network size and connectance (60 PERMANOVAs in total). We fit all PERMANOVAs using the R (R Core Team, 2016) function ‘adonis’ from the package *vegan* (Oksanen *et al.*, 2019) and calculated *p*-values using 9999 unstratified permutations. As conducting so many tests risks obtaining significant results by chance, we applied the correlated Bonferroni correction (Drezner & Drezner, 2016) before evaluating significance.

**Table 1:** Summary of analyses performed and results obtained therefrom. ‘S’ refers to species richness, ‘C’ connectance, ‘Ext’ to mean time to extinction across removals, ‘STL’ to shortest trophic level, and ‘Deg’ to degree (number of interaction partners).

| Question | Test | Result |
| --- | --- | --- |
| Are roles (count) related to Ext? | • PERMANOVA of Ext vs. role<br>• one per S-C combination (60 total) | • Yes. All PERMANOVA significant. |
| Are roles (count) similarly variable in species with long and short Ext? | • ANOVA of Disp(role) vs. Ext<br>• Disp(role) calculated by Betadisper<br>• one per S-C combination (60 total) | • No. All ANOVA significant. |
| Do species with long or short Ext have more variable roles? | • LM: Disp(role) vs. Ext | • Roles variability significantly increases with increasing Ext |
| Which motif counts, proportions, and Z-scores are most strongly related to Ext? (see Fig. 2). | • PLS: Ext vs. role, S, C, S:C, STL, Deg | • Ext decreases with omnivory but increases with other stable motifs. |
| How do motifs vary with Deg?<br>(see Fig. 3) | • LM: Count vs. Deg<br>• LM: Proportion vs. Deg<br>• LM: Z-score vs. Deg<br>• One LM per motif (39 total) | • All counts increase with increasing Deg<br>• Prop. omnivory increases with Deg, other stable motifs decrease<br>• All Z-scores increase with Deg |
| How do motifs vary with STL?<br>(see Fig. 3) | • LM: Count vs. STL<br>• LM: Proportion vs. STL<br>• LM: Z-score vs. STL<br>• One LM per motif (39 total) | • All counts increase with increasing STL<br>• Proportion of stable motifs except omnivory decrease with STL<br>• Z-scores of stable motifs decrease with STL |

In addition to participation in particular motifs affecting a species’ vulnerability to extinction, it may also be that those species most vulnerable to extinction (i.e. short time to extinction) have more variable roles than those which are least vulnerable, or vice versa. If there is unequal variance in roles across different times to extinction, this could cause false positive results in PERMANOVA tests. To account for this possibility, we first calculated the dispersion of roles for each value of mean time to extinction (treated categorically for this analysis), relative to the centroid for all species with the same mean time to extinction, using the function ‘betadisper’ from the R (R Core Team, 2016) package *vegan* (Oksanen *et al.*, 2019). We then tested whether some levels of mean time to extinction are associated with more widely-dispersed roles using an ANOVA test, fit using the function ‘anova’ within the package *vegan* (Oksanen *et al.*, 2019). Finally, to test whether role dispersion generally increases or decreases with increasing mean times to extinction, we fit a linear model relating role dispersion to mean time to extinction using the R (R Core Team, 2016) base function ‘lm’.

#### Identifying the motifs most strongly related to time to extinction

While the PERMANOVAs can tell us whether a species’ role as a whole is related to its mean time to extinction, it does not reveal which motifs have the strongest effect on time to extinction. To answer this question, we used a set of partial least squares (PLS) regressions to identify combinations of motifs which, together, explain substantial variation in time to extinction. Similar to a principal components analysis (PCA), PLS involves projecting the observed variables (in this case, participation in different motifs) into a new space and identifying latent variables made up of linear combinations of the observed variables (Mevik & Cederkvist, 2004; Mevik & Wehrens, 2007). In our case, these latent variables represent combinations of motifs which, together, explain substantial variation in mean time to extinction.

In the first PLS regression, we used mean time to extinction as the response and raw motif roles as well as network size, connectance, the interaction between size and connectance, in-degree (number of prey), and shortest trophic level (STL) (Hairston & Hairston, 1993) as predictors (Table 1). We include additional measures of network structure and species roles as these may affect the motif roles available to each focal species. To test whether any relationship between raw motif roles and time to extinction might be due to the tendency for high-degree species to appear in more motifs, we fit a second PLS regression using degree-normalized roles instead of the raw roles, with all other variables identical to the first regression. Finally, to understand whether it is the absolute frequency of motifs or the relative frequency compared to other species in the network that is related to time to extinction, we fit a third PLS regression using network-normalized roles as a predictor, with all other variables identical to the first regression.

We fit all regressions using the R (R Core Team, 2016) function ‘plsr’ from the package *pls* (Mevik & Wehrens, 2007). To prevent differences in range and intercept values from influencing the fit of the PLS model, we centered and scaled all variables. After initial fitting, we cross-validated each regression using 10 randomly-selected segments of the data and re-fitting the regression and then calculated the mean squared error of prediction (MSEP) for each model. MSEP is a measure of the error obtained when re-fitting a PLS or PCA model on test data, and is commonly used to select the optimum number of components (Mevik & Cederkvist, 2004). In order to balance obtaining a low MSEP with identifying a parsimonious model, we defined the optimum model as that with the fewest components that nevertheless had an MSEP within one standard deviation of the lowest MSEP obtained for any model. Model selection was performed using the R (R Core Team, 2016) function ‘selectNcomp’ from the package *plsr* (Mevik & Wehrens, 2007) using the method ‘onesigma’. After selecting the optimum number of components for each model, we re-fit the PLS regression including only the selected components. We then summed the coefficients of each predictor across axes to obtain an overall measure of the effect of each predictor on mean time to extinction.

#### Relating motif roles to other role definitions

Finally, we wanted to test the possibility that the relationships we observe between motif roles and time to extinction might be due to underlying relationships between species’ numbers of interaction partners (degree) or height within the food web (trophic level) and their motif roles. Species with more interaction partners will participate in more motifs (as each combination of three interacting species represents a unique motif). These species might also participate in a more diverse set of motifs, or might have roles in which certain motifs are over-represented. Similarly, species with very high (or very low) trophic levels may appear in fewer motifs than species with intermediate trophic levels, as they may not appear as often as prey (or predators) and therefore be excluded from some motifs.

To test these possibilities, we calculated each species’ degree (number of predators and prey) and trophic level. We defined trophic level as ‘shortest trophic level’ (STL): the length of the shortest food chain between the focal species and any basal resource (Hairston & Hairston, 1993). Basal resources are assigned a trophic level of one and other species are assigned an STL of one plus the STL of their prey. We then fit a series of linear regressions relating degree or STL and the count (taken from the raw motif role), proportion (taken from the degree-normalized motif role), and *Z*-score (taken from the network-normalized motif role) of each motif in the species’ role using a series of linear regressions (78 regressions total; Table 1). All regressions were fit with the R (R Core Team, 2016) base function ‘lm’. As with our PERMANOVA analysis, fitting so many regressions runs the risk of obtaining significant results by chance. To increase the likelihood of detecting a true relationship between species’ degrees and motif roles, we applied the correlated Bonferroni correction (Drezner & Drezner, 2016) before evaluating significance of these regressions.

We also wanted to estimate how the associations between motif roles and degree or STL might affect the relationships between motif roles and mean time to extinction. To do this, we fit two linear mixed-effect models relating mean time to extinction to a species’ degree (or STL). To account for differences in mean time to extinction across different network sizes and connectances, we included a random effect for the combination of network size and connectance. Both models were fit using the function ‘lmer’ from the R (R Core Team, 2016) package *lmerTest* (Kuznetsova *et al.*, 2017).

## Results

### Relating overall motif roles to mean time to extinction

Our series of PERMANOVA tests demonstrated that species’ overall raw motif roles were correlated with their mean time to extinction across all combinations of species richness and connectance (Table *S1, Appendix S2*). Taken individually, each PERMANOVA was significant (all *p*<0.025). Moreover, after applying the correlated Bonferroni correction (Drezner & Drezner, 2016), all PERMANOVAs remained significant. However, these significant results may be false positives since the variability of raw motif roles was not consistent across mean times to extinction for any combination of species richness and connectance (Table *S3, Appendix S2*). Specifically, longer mean times to extinction were associated with significantly more variable roles, and these relationships were stronger in smaller or less-connected networks. This means that there are many roles which are associated with long times to extinction but fewer roles which are associated with short times to extinction. After applying the correlated Bonferroni correction (Drezner & Drezner, 2016), both the ANOVAs testing for non-homogeneous variability of roles and regressions testing for relationships between role variability and time to extinction remained significant.

### Identifying the motifs most strongly related to mean time to extinction

#### Raw motif roles

The absolute number of times a species a species appeared in a motif was related to mean time to extinction, but not as strongly as trophic level and network properties. The optimum PLS model for mean time to extinction including raw motif roles included five components. The first two components explained 13.8% and 9.34% of variation in the mean time to extinction, with the remaining components explaining <1% of variation. Taking all components together, trophic level and the interaction between species richness and connectance had strong negative relationships with mean time to extinction while degree had the strongest positive relationship (Fig. 2A). The raw counts of the *omnivory* motif (S2) and several unstable motifs had negative relationships with mean time to extinction while the raw counts of the other stable motifs (S1, S4, and S5) as well as some unstable motifs had positive relationships with mean time to extinction. Although these effects appear small when considering the scaled, centered predictors, the large ranges of stable motifs which appear in species’ roles mean that the un-scaled effects of different numbers of motifs are actually large (Fig. 2B).

**Figure 2:**
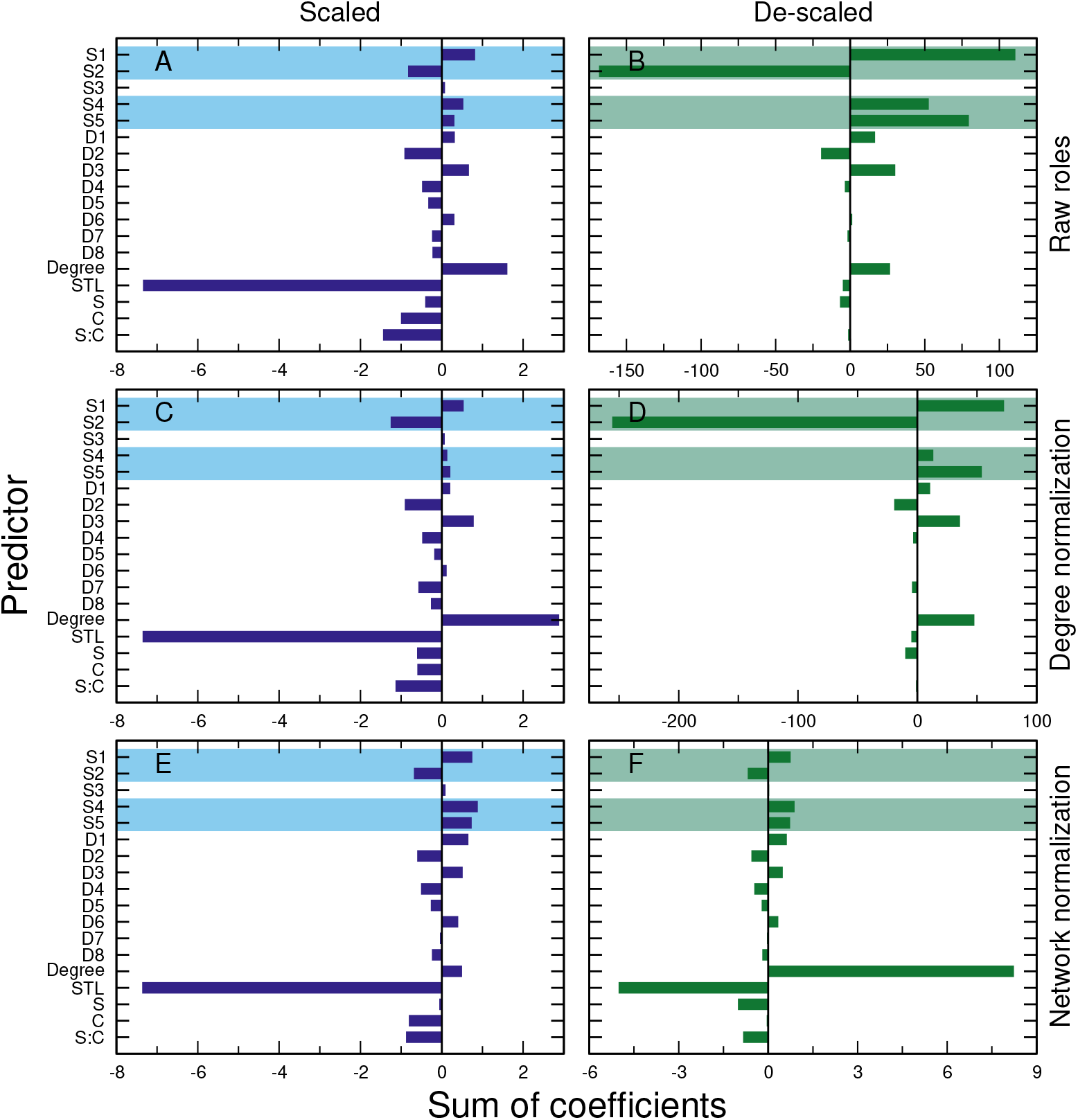
Sum of loadings of motif roles, degree, shortest trophic level (STL), web size (S), connectance (C), and their interaction (S:C) in the optimum number of axes in partial least squares regressions of mean time to extinction against all of the above predictors. We fit separate models for: **A-B)** raw motif counts, **C-D)** degree-normalized motif frequencies (i.e., counts of each motif divided by the total number of all motifs for the focal species) and **E-F)** network-normalized motif frequencies (i.e., *Z*-scores for each motif compared to all species in the network). Note that coefficients refer to centered and scaled predictors (i.e., an increase of one standard deviation in each predictor); stable motifs, degree, and species richness had much larger ranges in our dataset than the other predictors. Panels **A, C, E** (blue) give the sum of loadings for centered, scaled predictors. Panels **B, D, F** (green) give the sum of loadings after de-scaling predictors (i.e., for a unit increase in the scale of the predictor); stable motifs, degree, and species richness had much larger ranges in our dataset than the other predictors. In all panels, stable motifs are shaded. Motif names are as in Stouffer *et al.* (2007); see also Fig. S3, *Appendix S4*.

#### Degree-normalized roles

The proportions of a species’ role made up by different motifs showed stronger relationships to mean time to extinction than the raw motif frequencies. When considering degree-normalized motif roles, the optimum PLS model included five components. The first component explained 13.6% of variation in mean time to extinction, the second component explained 9.95% of variation, and the remaining components explained <1%. Taking all components together, trophic level had the strongest negative relationships with mean time to extinction while degree had the strongest positive relationship (Fig. 2C). The proportion of a species’ role made up by the *omnivory* motif had a similarly strong negative relationship to mean time to extinction to the interaction between species richness and connectance, while the proportions of a species’ role made up by the other stable motifs had weaker but positive relationships to mean time to extinction. The proportions of a species’ role made up of the unstable motifs again showed a mix of positive and negative associations with mean time to extinction.

#### Network-normalized roles

The *Z*-score of a species’ participation in each motif, relative to other species in the network, also showed a stronger relationship to mean time to extinction than raw motif roles. When considering network-normalized motif roles, the optimum PLS model included five components. The first component explained 18.8% of variation in mean time to extinction, the second component explained 3.32% of variance, the third component explained 1.78%, and the remaining components explained <1% of variation. Taking all components together, trophic level had the strongest negative relationships with mean time to extinction, while *Z*-scores for the stable *three-species chain* (S1), *direct competition* (S4), and *apparent competition* (S5) motifs had the strongest positive relationships with mean time to extinction (Fig. 2E). The *Z*-score of the stable *omnivory* motif had a similarly strong negative relationship to mean time to extinction as connectance and the interaction between species richness and connectance. The *Z*-scores of unstable motifs showed weaker positive and negative relationships to mean time to extinction.

### Relating motif roles to other role definitions

#### Degree

Species with higher degrees tended to have longer mean times to extinction (*β*=0.190, *p*<0.001). As expected, the count of all motifs increased significantly with increasing degree (Table *S3, Appendix S3*). The counts of the four stable motifs (S1, S2, S4, and S5) increased most rapidly with increasing degree (Fig. 3A). The proportions of a species’ role made up by the *three-species chain*, *direct competition*, and *apparent competition* motifs (S1, S4, and S5) in a species’ role decreased significantly with increasing degree, while the proportions of the *omnivory* (S2) and all unstable motifs increased significantly with increasing degree (Table *S3, Appendix S3*, Fig. 3C). The *Z*-scores of motifs relative to other species in the network all increased significantly with increasing degree (Table *S3, Appendix S3*). The *Z*-scores of the *omnivory* and *apparent competition* motifs (S2 and S5) increased most rapidly with degree, but there was not a clear separation in the slopes of stable and unstable motifs (Fig. 3E).

**Figure 3:**
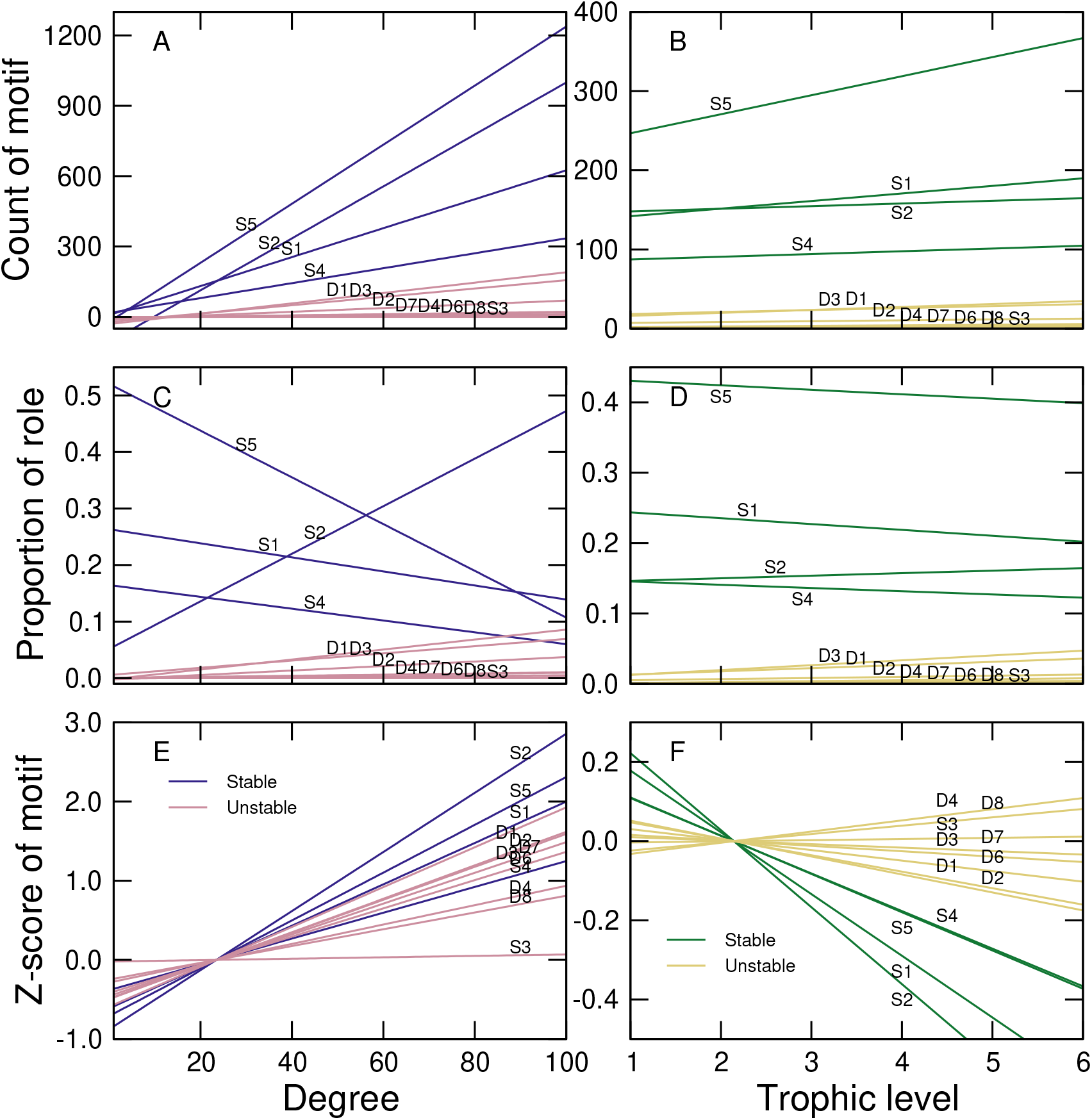
**A-B)** Species with more interaction partners or higher trophic levels partici-pated in more motifs. The counts of the four stable motifs (S1, S2, S4, and S5) increased most strongly with increasing degree and trophic level. **C-D)** The proportion of a species’ role made up by different motifs showed more variable trends. **C)** The proportion of the omnivory motif (S2) increased with increasing degree while those of the other stable motifs decreased. **D)** The proportions of all four stable motifs decreased with increasing trophic level. **E-F)** The *Z*-scores of all motifs increased with increasing degree, but trends were more variable with respect to trophic level. **F)** Species with high trophic levels tended to show under-representation of the four stable motifs while unstable motifs could be over- or under-represented. In all panels, motifs are plotted in order of the strength of their relationship with degree and all relationships were significant. Regressions for the four stable motifs are indicated by blue or green lines while unstable motifs are indicated by yellow or pink lines. 95% confidence intervals about the regression lines are too small to be visible in the current plot. Motif names are as in Stouffer *et al.* (2007); see also Fig. S3, *Appendix S4*.

#### Trophic level (height within food web)

Species with higher trophic levels tended to have shorter mean times to extinction (*β*=−11.0, *p*<0.001). The count of all motifs also increased with increasing trophic level (Table *S4, Appendix S3*). The counts of the four stable motifs (S1, S2, S4, and S5) were higher than counts of unstable motifs for all trophic levels, and the *apparent competition* (S5) and *three-species chain* (S1) also showed the most rapid increases with increasing trophic level (Fig. 3B). The proportions of a species’ role made up by the *three-species chain*, *direct competition*, and *apparent competition* (S1, S4, and S5) motifs decreased significantly with increasing trophic level, while the proportions of all other motifs increased significantly (Table *S4, Appendix S3; Supplemental Information*, Fig. 3D). The *Z*-scores of the four stable motifs, relative to other species in the network, decreased significantly with increasing trophic level while unstable motifs showed both significant increases and significant decreases with increasing trophic level (Table *S4, Appendix S3*, Fig. 3F).

## Discussion

We found that times to extinction were tightly correlated across removals. This suggests that the position of a focal species within a network can affect its risk of extinction following a disturbance regardless of where in the network the disturbance is applied. This result also justifies our use of mean times to extinction across removals as a measure of a species’ overall risk. Taken together, our results show that motif roles are related to mean times to extinction and that the omnivory motif may have a distinct effect from those of the other stable motifs, at least in body-mass structured vertebrate food webs. Testing whether these results hold true for other types of networks (e.g., invertebrate food-webs not structured by body mass) will require alternative simulation approaches; however, the increasing availability of highly-resolved empirical data for interactions among invertebrates () should allow researchers to create such simulations and test the generality of our results in the near future.

### Times to extinction are highly consistent

The consistency of species’ times to extinction across removals in our simulations suggests that some species are more likely to go extinct than others due to their position within a network. This complements other work identifying sets of species which are more vulnerable to extinction due to their traits (Curtsdotter *et al.*, 2011; Ryser *et al.*, 2019). Since the trophic level and degree of the species being removed can also affect which species, if any, go secondarily extinct (Wootton & Stouffer, 2016b; Dunne *et al.*, 2002), the properties of both disturbed and non-disturbed species affect the ways in which extinctions can cascade through a food web.

The stronger correlation of extinction orders in larger, more-connected networks, as well as the somewhat shorter mean times to extinction in these webs (Fig. S1, *Appendix S1*) may be due to the greater number of short pathways by which an extinction somewhere in the web can affect a focal species. These short pathways are more likely to have strong effects on the population dynamics of species along them than the longer indirect pathways in poorly-connected networks (Jordán & Scheuring, 2012; Jordán *et al.*, 2006). These stronger effects in turn likely explain why more secondary extinctions in large or highly-connected webs are due to indirect effects rather than direct loss of a prey or predator Wootton & Stouffer (2016b).

### Motif roles relate to extinction risk

Overall, our results show that species’ motif roles were related to their time to extinction after a disturbance (Table 1). Participating in more instances of the most stable motifs identified by Stouffer *et al.* (2007); Borrelli (2015), having a greater proportion of the role made up by these motifs, or appearing in significantly more of these motifs than other species in the network were all associated with longer times to extinction. Surprisingly, some unstable motifs were also associated with longer times to extinction. The unstable motifs are, however, rare in both empirical food webs (Stouffer *et al.*, 2007) and our simulated networks. Differences in participation in these motifs is therefore unlikely to have a large practical effect on species’ responses to a disturbance.

Part of the apparent benefit of participating in many stable motifs may be due to correlations between motif roles and degree. Counts of the stable motifs increased most rapidly with increasing degree, meaning that the relationship between high numbers of stable motifs and species persistence (Stouffer *et al.*, 2007; Borrelli, 2015) may be partly due to the beneficial effects of having a high degree (Cirtwill & Stouffer, 2016). That is, while a species which participates in many stable motifs is likely to benefit from damping of population cycles (Borrelli, 2015), it is also likely to maintain a reasonable number of food sources after a perturbation. The positive correlations between all four stable motifs and degree may also explain the greater variability among roles of species with long times to extinction. As species wither higher degrees also tend to participate in more motifs, there is a broader set of motif roles available to these species. Since high-degree species tend to participate in proportionally more stable than unstable motifs, many of the roles in this broad role-space should be associated with long times to extinction.

The relationships between trophic level and the counts or proportions of motifs tended to be weaker, although the *Z*-scores of all four stable motifs, tended to decrease strongly with increasing trophic level. Species with higher trophic levels also tended to have shorter mean times to extinction, suggesting that these species could be more susceptible due to being at the tops of longer food chains, participating in motifs that tend to amplify population fluctuations (Borrelli, 2015), or both. The correlations between degree, trophic level, and motif roles make identifying the precise mechanisms behind a species’ short or long mean time to extinction difficult. The possibility that these measures capture at least some independent information about the risk of secondary extinction, however, suggests that species with multiple ‘risky’ roles (i.e., low degree, participation in many unstable motifs, and/or high trophic level) may be more vulnerable than species with only one of the above.

### Omnivory may be an especially informative motif

Although there may be many motif roles which promote long times to extinction, not all of the stable motifs are equally likely to be included in such roles. While the three-species chain, apparent competition, and direct competition motifs were all associated with longer times to extinction as expected, the omnivory motif was associated with shorter mean times to extinction. This is consistent with an omnivory motif being less likely to retain all three species than the chain, apparent competition, or direct competition motifs when modeled in isolation (Borrelli, 2015).

It may be difficult to isolate the effect of omnivory, however, as the proportion of the omnivory motif in a species’ role increased strongly with increasing degree. This relationship could partly explain the variety of earlier work showing that the omnivory can be stable or unstable under different conditions and may or may not increase the overall stability of an entire food web (McCann & Hastings, 1997; Emmerson & Yearsley, 2004; Borrelli, 2015; Monteiro & Faria, 2016). If high-degree species tend to persist for a long time but high-omnivory species are more likely to go extinct, then it may be difficult to predict the outcome for a species with both high degree and a role rich in the omnivory motif. However, this relationship does suggest that motif roles may help us to compare risk for species with similar degrees.

### Moving forward with motifs

As motif roles become more common as tools for ecologists wishing to understand how species’ positions within networks are related to their traits (Cirtwill & Eklöf, 2018), taxonomy (Stouffer *et al.*, 2007), and position in space or time (Baker *et al.*, 2015), it is natural to wonder how these roles might relate to a species’ response to disturbance. Our results suggest that participation in the three-species chain, apparent competition, and direct competition motifs promote longer persistence after a disturbance, while the omnivory motif may indicate more rapid extinction. However, interpreting these trends is complicated by strong relationships between motif roles and degree, which itself promotes longer persistence after a disturbance.

Because relationships between motif roles and degree appear even when considering the proportion of species roles made up by each motif, as in Baker *et al.* (2015); Cirtwill & Stouffer (2015); Simmons *et al.* (2019), normalizing motif roles based on the total number of motifs does not control for differences in species’ roles due to degree. The papers cited above explicitly aim to control for differences due to the total number of motifs in which a species appears (which depends upon the degrees of the focal species *and* those of its interaction partners) rather than degree *per se*. However, this distinction is subtle (hence our use of ‘degree-normalization’ to describe the proportional roles) and future authors should take great care that they are not using the sum of motifs as a substitute for degree as our results show that these quantities are not interchangeable. Rather, our results suggest that both motif roles and simpler measures of network structure can provide information about a species’ response to disturbance. Motifs may be particularly useful when comparing species with similar or trophic levels: the multi-dimensional nature of the motif role allows it to capture information which is lost in single-value measures such as degree or trophic level. While the interpretation of motif roles remains challenging, the relationships we have found between these roles and time to extinction suggest that they are worth retaining in the toolbox of ecological network analysis.

## Acknowledgments

We thank Eva Delmas and Chris Rackauckas for their kind assistance with troubleshooting the simulation model and the Spatial Foodweb Ecology Group for providing feedback on the manuscript. ARC is supported by a Finnish Academy Postdoctoral research grant (#332999).

## Appendices

All of the following Appendices are included as supplemental information.

## S1 Testing consistence of times to extinction

Supplemental methods and results related to testing whether the identity of the removed species has a large effect on the time to extinction for non-removed species. Includes Figure S1.

## S2 Details of PERMANOVA results

Supplemental results for PERMANOVAs testing whether species’ overall roles are related to their mean times to extinction. Includes **Figure S2**, **Tables S1-S2**.

## S3 Relating motifs roles to other measures

Gives the slopes of relationships between motif roles and degree or shortest trophic level (STL) in **Tables S3-S4**.

## S4 Motif labels

**Figure S3** illustrates the 13 unique three-species motifs and labels each one according to the naming scheme in Stouffer *et al.* (2007).

