## Appendices S1-S3 for "Stable motifs delay species loss in simulated food webs"

#### Table of Contents

##### S1: Testing consistence of times to extinction

Supplemental methods and results related to testing whether the identity of the removed species has a large effect on the time to extinction for non-removed species. Includes **Figure S1**.

##### S2: Details of PERMANOVA results

Supplemental results for PERMANOVAs testing whether species' overall roles are related to their mean times to extinction. Includes **Figure S2, Tables S1-S2**.

##### S3: Relating motifs roles to other measures

Gives the slopes of relationships between motif roles and degree or shortest trophic level (STL) in **Tables S3-S4**.

##### S4: Motif labels

**Figure S3** illustrates the 13 unique three-species motifs and labels each one according to the naming scheme in Stouffer *et al.* (2007).

### S1: Testing consistency of times to extinction across removals

#### Methods

We simulated population dynamics within each food web after removing each species separately. In order to assess species' overall vulnerability to the loss of an arbitrary other species within the food web, we used mean time to extinction across all removals as a response. To ensure that this is a robust measure, we calculated the Pearson correlation of times to extinction for each species across all extinctions in each network. We then tested whether the strength of these correlations varied with species richness and/or connectance by fitting a general linear model including fixed effects of species richness, connectance, and their interaction, as well as a random effect for network ID. We fit the model using the R (R Core Team, 2016) function 'lmer' from the package *lmerTest* (Kuznetsova *et al.*, 2017).

#### Results

In general, time to extinction was highly correlated across removals (Fig. S1). The mean Pearson correlation for times to extinction across all removals within a network was 0.903 (range: 0.512-0.973). This means that, in general, the species which go extinct fastest after species $i$ is removed also go extinct fastest after species $j$ is removed. The correlation was stronger in larger webs, particularly those with high connectance ($\beta_S$=1.33×10$^-$3, $p$<0.001; $\beta_C$=-8.08×10$^-$2, $p$<0.001; and $\beta_{S:C}$=2.12×10$^-$3, $p$<0.001, respectively). Mean time to extinction is, therefore, a good measure of a species' overall vulnerability.

**Figure S1: A)** Time to extinction for each species within a simulated network was highly correlated across removals. Circles indicate the mean correlation of time to extinction across removals for all species in all 100 simulated networks for a given combination of species richness and connectance. Lines indicate the predicted correlation based on the fixed effects of a linear model including species richness, connectance, and the interaction between them, as well as a random effect of network. Symbol and line colors indicate connectance. **B)** Mean time to extinction across all species within a network was slightly longer in small and less-connected networks. As in **A)**, line colors indicate connectance. **C)** Mean time to extinction was more strongly associated with connectance than species richness, with more-connected networks having shorter mean times to extinction. Line colors indicate species richness.

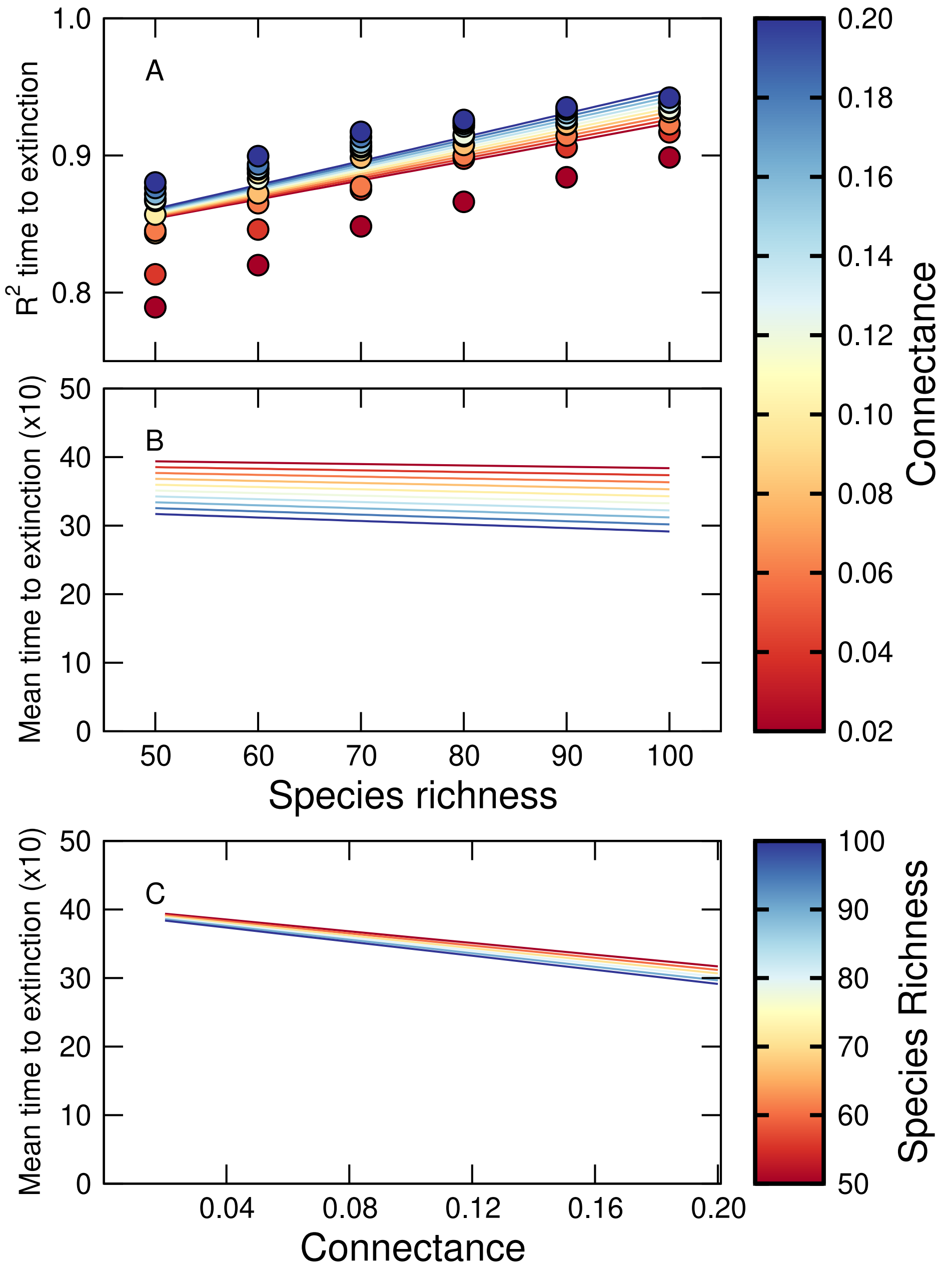

#### S2: Details of PERMANOVA results

All PERMANOVAs were significant, indicating that species' overall motif roles are related to their mean times to extinction.

**Figure S2:** Here we show (**A**) the pseudo-$F$ statistics and (**B**) the $p$-values for each PERMANOVA relating species' roles to their mean extinction order when all species in the web are separately removed. We fit one PERMANOVA per combination of species richness and connectance. $p$-values for each PERMANOVA are based on 9999 permutations, stratified by network. Symbols below the dotted line in **B** indicate a significant $p$-values.

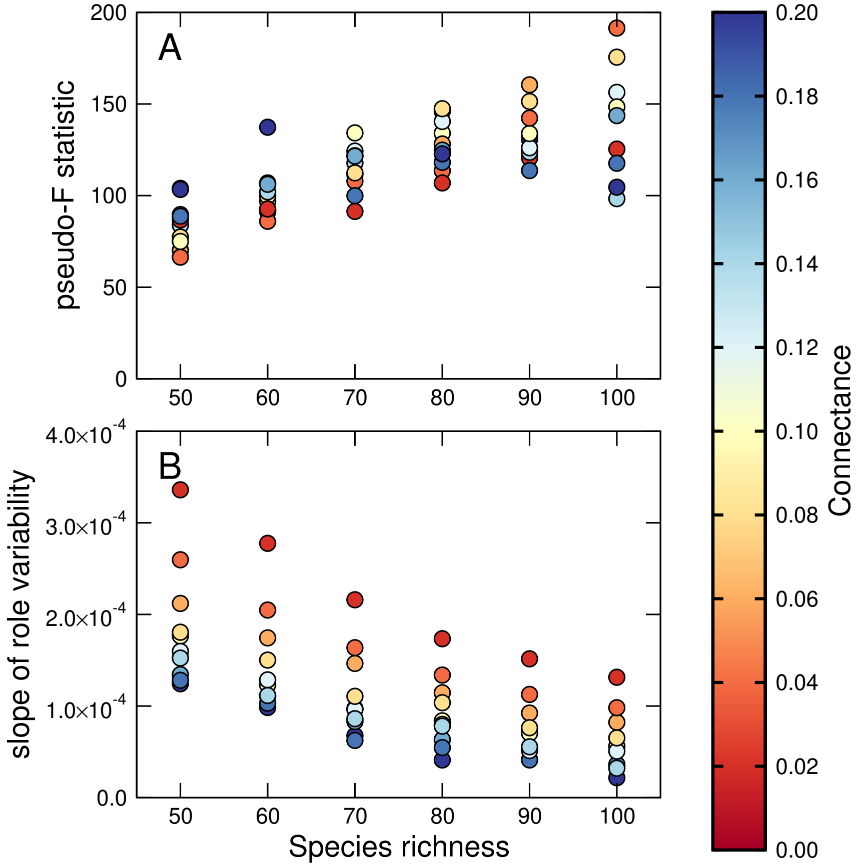

**Table S1:** For each combination of species richness (S) and connectance (C), the mean extinction order of a focal species was related to its raw motif role. We tested this using a series of PERMANOVAs with 9999 permutations each. Here we show the mean correlation among extinction orders across all removed species ($R^2$) and all 100 simulated networks for each combination of S and C, as well as the pseudo-$F$ statistic and $p$-value for each PERMANOVA. All tests remained significant after applying the correlated Bonferroni correction (Drezner & Drezner, 2016).

| S | C | $R^2$ | pseudo-$F$ | $p$-value | S | C | $R^2$ | pseudo-$F$ | $p$-value |
|---|---|---|---|---|---|---|---|---|---|
| 50 | 0.02 | 0.789 | 86.7 | 0.017 | 80 | 0.02 | 0.866 | 107 | 0.013 |
| 50 | 0.04 | 0.813 | 66.4 | 0.013 | 80 | 0.04 | 0.898 | 114 | 0.014 |
| 50 | 0.06 | 0.845 | 70.2 | 0.014 | 80 | 0.06 | 0.9 | 128 | 0.016 |
| 50 | 0.08 | 0.843 | 77.4 | 0.015 | 80 | 0.08 | 0.908 | 147 | 0.018 |
| 50 | 0.10 | 0.857 | 75.0 | 0.015 | 80 | 0.10 | 0.914 | 134 | 0.016 |
| 50 | 0.12 | 0.868 | 104 | 0.020 | 80 | 0.12 | 0.915 | 140 | 0.017 |
| 50 | 0.14 | 0.867 | 83.8 | 0.016 | 80 | 0.14 | 0.921 | 146 | 0.018 |
| 50 | 0.16 | 0.872 | 89.7 | 0.018 | 80 | 0.16 | 0.923 | 125 | 0.015 |
| 50 | 0.18 | 0.876 | 88.7 | 0.017 | 80 | 0.18 | 0.925 | 118 | 0.015 |
| 50 | 0.20 | 0.88 | 103 | 0.020 | 80 | 0.20 | 0.926 | 123 | 0.015 |
| 60 | 0.02 | 0.82 | 92.7 | 0.015 | 90 | 0.02 | 0.884 | 121 | 0.013 |
| 60 | 0.04 | 0.846 | 86.0 | 0.014 | 90 | 0.04 | 0.906 | 142 | 0.016 |
| 60 | 0.06 | 0.865 | 91.2 | 0.015 | 90 | 0.06 | 0.915 | 160 | 0.018 |
| 60 | 0.08 | 0.872 | 99.7 | 0.016 | 90 | 0.08 | 0.923 | 151 | 0.017 |
| 60 | 0.10 | 0.887 | 92.1 | 0.015 | 90 | 0.10 | 0.923 | 128 | 0.014 |
| 60 | 0.12 | 0.883 | 96.9 | 0.016 | 90 | 0.12 | 0.927 | 128 | 0.014 |
| 60 | 0.14 | 0.891 | 102 | 0.017 | 90 | 0.14 | 0.928 | 126 | 0.014 |
| 60 | 0.16 | 0.89 | 106 | 0.017 | 90 | 0.16 | 0.931 | 138 | 0.015 |
| 60 | 0.18 | 0.893 | 107 | 0.018 | 90 | 0.18 | 0.934 | 107 | 0.012 |
| 60 | 0.20 | 0.899 | 137 | 0.022 | 90 | 0.20 | 0.936 | 127 | 0.014 |
| 70 | 0.02 | 0.848 | 91.4 | 0.013 | 100 | 0.02 | 0.899 | 125 | 0.012 |
| 70 | 0.04 | 0.875 | 108 | 0.015 | 100 | 0.04 | 0.917 | 191 | 0.019 |
| 70 | 0.06 | 0.877 | 111 | 0.016 | 100 | 0.06 | 0.923 | 206 | 0.020 |
| 70 | 0.08 | 0.898 | 112 | 0.016 | 100 | 0.08 | 0.932 | 176 | 0.017 |
| 70 | 0.10 | 0.904 | 134 | 0.019 | 100 | 0.10 | 0.934 | 148 | 0.015 |
| 70 | 0.12 | 0.907 | 124 | 0.017 | 100 | 0.12 | 0.934 | 156 | 0.015 |
| 70 | 0.14 | 0.906 | 118 | 0.017 | 100 | 0.14 | 0.939 | 98.3 | 0.010 |
| 70 | 0.16 | 0.909 | 122 | 0.017 | 100 | 0.16 | 0.938 | 144 | 0.014 |
| 70 | 0.18 | 0.913 | 99.9 | 0.014 | 100 | 0.18 | 0.939 | 118 | 0.012 |
| 70 | 0.20 | 0.917 | 122 | 0.017 | 100 | 0.20 | 0.942 | 105 | 0.010 |

**Table S2:** For each combination of species richness (S) and connectance (C), some levels of mean time to extinction were associated with more variable roles than others. This may cause false positives in the PERMANOVAs reported in table S1. In all cases, the variability of species' roles increased significantly with increasingly long mean times to extinction. All tests remained significant after applying the correlated Bonferroni correction (Drezner & Drezner, 2016).

| | | ANOVA | | Regression | | | | ANOVA | | Regression | |
|---|---|---|---|---|---|---|---|---|---|---|---|
| S | C | $F$ | $p$-value | $\beta$ | $p$-value | S | C | $F$ | $p$-value | $\beta$ | $p$-value |
| 50 | 0.02 | 3.38 | <0.001 | $3.36\times10^{-4}$ | <0.001 | 80 | 0.02 | 3.72 | <0.001 | $1.74\times10^{-4}$ | <0.001 |
| 50 | 0.04 | 3.90 | <0.001 | $2.60\times10^{-4}$ | <0.001 | 80 | 0.04 | 3.87 | <0.001 | $1.34\times10^{-4}$ | <0.001 |
| 50 | 0.06 | 3.75 | <0.001 | $2.12\times10^{-4}$ | <0.001 | 80 | 0.06 | 3.86 | <0.001 | $1.15\times10^{-4}$ | <0.001 |
| 50 | 0.08 | 3.59 | <0.001 | $1.81\times10^{-4}$ | <0.001 | 80 | 0.08 | 3.83 | <0.001 | $1.04\times10^{-4}$ | <0.001 |
| 50 | 0.10 | 3.56 | <0.001 | $1.76\times10^{-4}$ | <0.001 | 80 | 0.10 | 3.59 | <0.001 | $8.39\times10^{-5}$ | <0.001 |
| 50 | 0.12 | 3.76 | <0.001 | $1.60\times10^{-4}$ | <0.001 | 80 | 0.12 | 3.49 | <0.001 | $8.00\times10^{-5}$ | <0.001 |
| 50 | 0.14 | 3.39 | <0.001 | $1.52\times10^{-4}$ | <0.001 | 80 | 0.14 | 3.27 | <0.001 | $7.80\times10^{-5}$ | <0.001 |
| 50 | 0.16 | 3.34 | <0.001 | $1.34\times10^{-4}$ | <0.001 | 80 | 0.16 | 3.02 | <0.001 | $6.34\times10^{-5}$ | <0.001 |
| 50 | 0.18 | 3.23 | <0.001 | $1.28\times10^{-4}$ | <0.001 | 80 | 0.18 | 2.72 | <0.001 | $5.46\times10^{-5}$ | <0.001 |
| 50 | 0.20 | 3.12 | <0.001 | $1.24\times10^{-4}$ | <0.001 | 80 | 0.20 | 2.55 | <0.001 | $4.11\times10^{-5}$ | <0.001 |
| 60 | 0.02 | 3.58 | <0.001 | $2.78\times10^{-4}$ | <0.001 | 90 | 0.02 | 3.56 | <0.001 | $1.52\times10^{-4}$ | <0.001 |
| 60 | 0.04 | 3.88 | <0.001 | $2.05\times10^{-4}$ | <0.001 | 90 | 0.04 | 4.00 | <0.001 | $1.13\times10^{-4}$ | <0.001 |
| 60 | 0.06 | 3.72 | <0.001 | $1.74\times10^{-4}$ | <0.001 | 90 | 0.06 | 3.64 | <0.001 | $9.23\times10^{-5}$ | <0.001 |
| 60 | 0.08 | 3.47 | <0.001 | $1.50\times10^{-4}$ | <0.001 | 90 | 0.08 | 3.44 | <0.001 | $7.63\times10^{-5}$ | <0.001 |
| 60 | 0.10 | 3.58 | <0.001 | $1.23\times10^{-4}$ | <0.001 | 90 | 0.10 | 3.24 | <0.001 | $7.03\times10^{-5}$ | <0.001 |
| 60 | 0.12 | 3.45 | <0.001 | $1.29\times10^{-4}$ | <0.001 | 90 | 0.12 | 3.02 | <0.001 | $5.14\times10^{-5}$ | <0.001 |
| 60 | 0.14 | 3.45 | <0.001 | $1.12\times10^{-4}$ | <0.001 | 90 | 0.14 | 2.85 | <0.001 | $5.56\times10^{-5}$ | <0.001 |
| 60 | 0.16 | 3.30 | <0.001 | $1.11\times10^{-4}$ | <0.001 | 90 | 0.16 | 2.92 | <0.001 | $5.26\times10^{-5}$ | <0.001 |
| 60 | 0.18 | 3.07 | <0.001 | $1.03\times10^{-4}$ | <0.001 | 90 | 0.18 | 2.58 | <0.001 | $4.11\times10^{-5}$ | <0.001 |
| 60 | 0.20 | 3.15 | <0.001 | $9.84\times10^{-5}$ | <0.001 | 90 | 0.20 | 2.59 | <0.001 | $4.17\times10^{-5}$ | <0.001 |
| 70 | 0.02 | 3.45 | <0.001 | $2.16\times10^{-4}$ | <0.001 | 100 | 0.02 | 3.99 | <0.001 | $1.31\times10^{-4}$ | <0.001 |
| 70 | 0.04 | 3.76 | <0.001 | $1.64\times10^{-4}$ | <0.001 | 100 | 0.04 | 3.95 | <0.001 | $9.81\times10^{-5}$ | <0.001 |
| 70 | 0.06 | 3.83 | <0.001 | $1.46\times10^{-4}$ | <0.001 | 100 | 0.06 | 3.65 | <0.001 | $8.23\times10^{-5}$ | <0.001 |
| 70 | 0.08 | 3.64 | <0.001 | $1.11\times10^{-4}$ | <0.001 | 100 | 0.08 | 3.31 | <0.001 | $6.52\times10^{-5}$ | <0.001 |
| 70 | 0.10 | 3.39 | <0.001 | $9.72\times10^{-5}$ | <0.001 | 100 | 0.10 | 3.11 | <0.001 | $5.62\times10^{-5}$ | <0.001 |
| 70 | 0.12 | 3.42 | <0.001 | $9.64\times10^{-5}$ | <0.001 | 100 | 0.12 | 3.05 | <0.001 | $5.07\times10^{-5}$ | <0.001 |
| 70 | 0.14 | 3.09 | <0.001 | $8.62\times10^{-5}$ | <0.001 | 100 | 0.14 | 2.54 | <0.001 | $3.22\times10^{-5}$ | <0.001 |
| 70 | 0.16 | 3.05 | <0.001 | $8.31\times10^{-5}$ | <0.001 | 100 | 0.16 | 2.56 | <0.001 | $3.44\times10^{-5}$ | <0.001 |
| 70 | 0.18 | 2.85 | <0.001 | $6.25\times10^{-5}$ | <0.001 | 100 | 0.18 | 2.58 | <0.001 | $3.63\times10^{-5}$ | <0.001 |
| 70 | 0.20 | 2.81 | <0.001 | $6.77\times10^{-5}$ | <0.001 | 100 | 0.20 | 2.32 | <0.001 | $2.17\times10^{-5}$ | <0.001 |

### S3: Relating motif roles to other measures

**Table S3:** Slopes and $p$-values for linear regressions relating degree (number of interaction partners) to the raw counts of motifs, degree-normalized proportions of motifs, and network-normalized $Z$-scores of motifs. Motifs are named as in Stouffer *et al.* (2007). Motifs S1 (*three-species chain*), S2 (*omnivory*), S4 (*direct competition*), and S5 (*apparent competition*) are the 'stable' motifs of particular interest in our study (highlighted in bold). All regressions remained significant after applying the correlated Bonferroni correction (Drezner & Drezner, 2016).

| Motif | Raw roles | | Degree-normalized roles | | Network-normalized roles | |
|---|---|---|---|---|---|---|
| | Slope | $p$-value | Slope | $p$-value | Slope | $p$-value |
| **S1** | 6.17 | **<0.001** | $-1.24\times10^{-3}$ | **<0.001** | $2.61\times10^{-2}$ | **<0.001** |
| **S2** | 11.1 | **<0.001** | $4.21\times10^{-3}$ | **<0.001** | $3.73\times10^{-2}$ | **<0.001** |
| **S4** | 3.18 | **<0.001** | $-1.04\times10^{-3}$ | **<0.001** | $1.63\times10^{-2}$ | **<0.001** |
| **S5** | 12.6 | **<0.001** | $-4.13\times10^{-3}$ | **<0.001** | $3.02\times10^{-2}$ | **<0.001** |
| S3 | $6.73\times10^{-4}$ | **<0.001** | $2.96\times10^{-7}$ | **<0.001** | $8.90\times10^{-4}$ | **<0.001** |
| D1 | 2.21 | **<0.001** | $8.83\times10^{-4}$ | **<0.001** | $2.52\times10^{-2}$ | **<0.001** |
| D2 | 0.797 | **<0.001** | $3.98\times10^{-4}$ | **<0.001** | $2.11\times10^{-2}$ | **<0.001** |
| D3 | 1.78 | **<0.001** | $6.37\times10^{-4}$ | **<0.001** | $2.08\times10^{-2}$ | **<0.001** |
| D4 | 0.203 | **<0.001** | $8.98\times10^{-5}$ | **<0.001** | $1.22\times10^{-2}$ | **<0.001** |
| D5 | $3.77\times10^{-2}$ | **<0.001** | $1.52\times10^{-5}$ | **<0.001** | $1.11\times10^{-2}$ | **<0.001** |
| D6 | 0.127 | **<0.001** | $5.85\times10^{-5}$ | **<0.001** | $1.78\times10^{-2}$ | **<0.001** |
| D7 | 0.256 | **<0.001** | $1.15\times10^{-4}$ | **<0.001** | $1.94\times10^{-2}$ | **<0.001** |
| D8 | $4.04\times10^{-2}$ | **<0.001** | $1.61\times10^{-5}$ | **<0.001** | $1.06\times10^{-2}$ | **<0.001** |

**Table S4:** Slopes and $p$-values for linear regressions relating trophic level (STL; height in the food web) to the raw counts of motifs, degree-normalized proportions of motifs, and network-normalized $Z$-scores of motifs. Motifs are named as in Stouffer *et al.* (2007). Motifs S1 (*three-species chain*), S2 (*omnivory*), S4 (*direct competition*), and S5 (*apparent competition*) are the 'stable' motifs of particular interest in our study (highlighted in bold). All regressions remained significant after applying the correlated Bonferroni correction (Drezner & Drezner, 2016).

| Motif | Raw roles | | Degree-normalized roles | | Network-normalized roles | |
|---|---|---|---|---|---|---|
| | Slope | $p$-value | Slope | $p$-value | Slope | $p$-value |
| **S1** | 9.58 | **<0.001** | -8.32×$10^{-3}$ | **<0.001** | -0.156 | **<0.001** |
| **S2** | 3.36 | **<0.001** | 3.62×$10^{-3}$ | **<0.001** | -0.194 | **<0.001** |
| **S4** | 3.48 | **<0.001** | -4.59×$10^{-3}$ | **<0.001** | -9.51×$10^{-2}$ | **<0.001** |
| **S5** | 24.0 | **<0.001** | -6.35×$10^{-3}$ | **<0.001** | -9.66×$10^{-2}$ | **<0.001** |
| S3 | 2.96×$10^{-3}$ | **<0.001** | 4.82×$10^{-6}$ | **<0.001** | 2.91×$10^{-3}$ | **<0.001** |
| D1 | 2.48 | **<0.001** | 4.50×$10^{-3}$ | **<0.001** | -4.16×$10^{-2}$ | **<0.001** |
| D2 | 1.09 | **<0.001** | 1.61×$10^{-3}$ | **<0.001** | -4.53×$10^{-2}$ | **<0.001** |
| D3 | 3.72 | **<0.001** | 6.89×$10^{-3}$ | **<0.001** | -1.37×$10^{-2}$ | **<0.001** |
| D4 | 0.804 | **<0.001** | 1.34×$10^{-3}$ | **<0.001** | 2.82×$10^{-2}$ | **<0.001** |
| D5 | 0.112 | **<0.001** | 2.05×$10^{-4}$ | **<0.001** | 1.85×$10^{-2}$ | **<0.001** |
| D6 | 9.93×$10^{-2}$ | **<0.001** | 1.86×$10^{-4}$ | **<0.001** | -2.65×$10^{-2}$ | **<0.001** |
| D7 | 0.380 | **<0.001** | 6.64×$10^{-4}$ | **<0.001** | -8.81×$10^{-3}$ | **<0.001** |
| D8 | 0.135 | **<0.001** | 2.46×$10^{-4}$ | **<0.001** | 2.10×$10^{-2}$ | **<0.001** |

#### S4: Motif labels

**Figure S3:** There are 13 unique three-species motifs which can appear in food webs. Motifs S1, S2, S4, and S5 have been identified as more stable than other motifs when modeled in isolation and as potentially increasing the stability of empirical food webs. Labels are as in Stouffer *et al.* (2007).

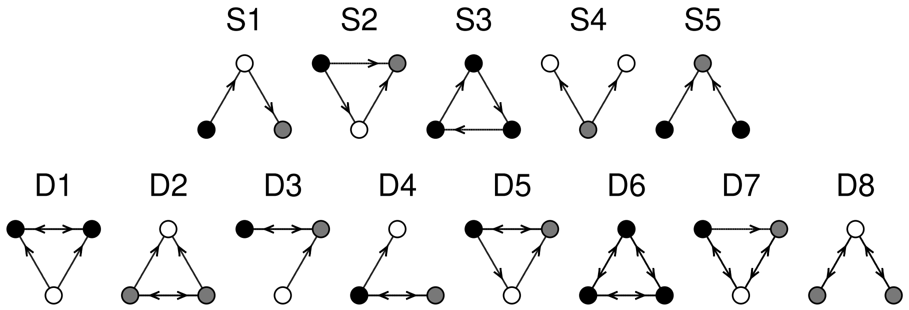


S1: Three-species chain
S2: Omnivory
S3: One-way three-species loop
S4: Direct competition
S5: Apparent competition
D1: S4 with mutual predation among predators
D2: S5 with mutual predation among prey
D3: S1 with mutual predation among top and intermediate spp.
D4: S1 with mutual predation among bottom and intermediate spp.
D5: S2 with mutual predation among top and bottom spp.
D6: Two-way three-species loop
D7: Three-species loop with two two-way links
D8: S1 with mutual predation along both links.
